# Impact of Sequential Processing (Boiling and Fermentation) on the Nutritional, Anti-nutrient, and Antioxidant Profile of *Manihot esculenta* Tubers

**DOI:** 10.64898/2026.03.03.709334

**Authors:** Grace Edet Bassey, Etukudo Okon Jimmy, Helen Titilope Olatunbosun

## Abstract

1.

**Background:** *Manihot esculenta* (Cassava) is a vital staple in Sub-Saharan Africa, yet its high levels of **cyanogenic glycosides** and **anti-nutrients** pose health risks. While boiling is common, its holistic impact on the **nutritional biochemistry** and **antioxidant** profile of the “Farmer’s Pride” (IBA 961632) variety remains under-characterized. This study evaluated the sequential impact of **food processing** –boiling and multi-stage **fermentation –**on cassava’s toxicological and bioactive profiles.

**Methods:** Fresh tubers were boiled for 10 minutes and fermented for 24, 48, and 72 hours. Proximate composition, vitamins, and anti-nutritional factors (cyanide, oxalate, phytate) were quantified. **Linamarase** activity and total phenolic and flavonoid contents were measured to assess enzymatic detoxification and phytotherapeutic potential.

**Results:** Boiling concentrated carbohydrates but created a “nutrient void,” leaching 93% of Vitamin C. However, fermentation acted as a biochemical refinery; by 72 hours, total cyanide plummeted from 98.15 to 0.54 mg/100g, meeting WHO safety standards. Concurrently, fermentation triggered a resurgence in bioactives, significantly increasing phenolic and flavonoid levels.

**Conclusion:** Boiling alone is insufficient for detoxification. Sequential fermentation beyond 48 hours is essential to “rescue” **antioxidant** potential and ensure safety. The 72-hour fermented tuber represents an optimized **bioactive food vehicle** for managing oxidative stress-related pathologies like **prostatic hyperplasia**

## 2. Introduction

Cassava (*Manihot esculenta* Crantz) serves as a vital staple crop for over 800 million people globally, particularly in sub-Saharan Africa, due to its resilience to drought and high carbohydrate yield [1]. Beyond its caloric value, cassava tubers contain essential micronutrients, including Vitamin C, Vitamin E, and various minerals, which are critical for the nutritional security of low-income populations [2, 3]. However, the utilization of cassava is significantly hindered by the presence of endogenous cyanogenic glycosides, primarily linamarin and lotaustralin.

Upon tissue damage, these compounds are hydrolyzed by the enzyme linamarase to release hydrogen cyanide (HCN), a potent toxin that can cause acute poisoning or chronic neurological disorders such as konzo and tropical ataxic neuropathy if consumed in inadequately processed forms [4].

In addition to cyanogens, cassava tubers contain other anti-nutritional factors (ANFs) such as oxalates and phytates, which impede the bioavailability of essential minerals like calcium, zinc, and iron [5]. High levels of these antagonists are associated with micronutrient deficiencies and stunted growth in communities where cassava is the primary dietary component. Consequently, effective processing techniques are mandatory to reduce these toxins to safe levels. The World Health Organization (WHO) and the Food and Agriculture Organization (FAO) of the United Nations have established a safety threshold of 10 mg/kg (10 ppm) for total hydrocyanic acid in cassava flour intended for human consumption [6].

Common processing methods, such as boiling, are effective at reducing initial cyanide levels and softening the tuber for consumption. However, boiling often leads to a significant “nutrient trade-off,” where water-soluble vitamins and antioxidant phenolics are lost through thermal degradation and leaching into the processing water [7]. To mitigate these losses and further detoxify the tuber, fermentation is frequently employed. Fermentation not only facilitates the near-complete removal of cyanogens through microbial enzymatic activity but also enhances the nutritional profile by releasing bound bioactive compounds and potentially synthesizing new vitamins through the action of lactic acid bacteria and yeasts [8].

Recent studies have highlighted the role of fermentation time in modulating the antioxidant capacity of cassava [9]. During fermentation, microbial beta-glucosidases break down complex plant cell wall structures, leading to a progressive increase in total phenolic content (TPC) and total flavonoid content (TFC), thereby improving the therapeutic potential of the food vehicle [10]. Despite the known benefits of individual processing steps, there is limited data on the sequential impact of boiling followed by multi-day fermentation on the holistic biochemical profile, including vitamins, antioxidants, and a wide array of anti-nutrients, of Nigerian cassava varieties.

Therefore, this study aimed to characterize the biochemical transformations that occur in *M. esculenta* tubers during sequential processing. Specifically, we evaluated how boiling and subsequent fermentation stages (24 to 72 hours) influence proximate composition, vitamin retention, and antioxidant capacity, while monitoring the reduction of diverse anti-nutritional factors. These findings are essential for optimizing cassava processing to produce safe, nutrient-dense products suitable for therapeutic applications.

## 3. Materials and Methods

### 3.1 Plant Collection and Identification

Fresh cassava (*Manihot esculenta* Crantz, Family: Euphorbiaceae) tubers of the **Farmer’s Pride (IBA 961632)** variety were harvested from a local farm in **Ikot Odiong Ididep**, Akwa Ibom State, Nigeria. The plant material was formally identified and authenticated by the Department of Botany, University of Uyo. Further botanical validation and variety coding were provided by the **Cassava Program, National Root Crops Research Institute (NRCRI), Umudike**, Abia State. A voucher specimen was deposited at the University of Uyo herbarium for future reference.

### 3.2 Sample Preparation and Processing

The tubers were peeled, washed with deionized water, and sliced into uniform chips. The samples were divided into five treatment groups based on processing stages:

- **Fresh:** Raw, unprocessed cassava.
- **Boiled:** Tubers were submerged in boiling water and cooked for **10 minutes** using a camp gas stove. The pot was left **uncovered** throughout the process to facilitate the volatilization of hydrogen cyanide.
- **D1 (24h Fermented):** Boiled tubers were submerged in distilled water in sterile containers and allowed to ferment at room temperature (approx. 28 +/- 2°C) for 24 hours.
- **D2 (48h Fermented):** Boiled tubers fermented for 48 hours.
- **D3 (72h Fermented):** Boiled tubers fermented for 72 hours.

Immediately following each processing stage, the samples were drained and stored in sterile, airtight containers under refrigeration (4°C) to arrest further fermentation and enzymatic activity. All biochemical and spectrophotometric analyses were conducted on these refrigerated samples within a timeframe that ensured maximum nutrient retention.

### 3.3 Determination of Proximate Composition

Proximate analysis, including moisture content, crude protein (Kjeldahl method), crude lipid (Soxhlet extraction), crude fiber, and ash content, was conducted following the standard procedures of the **Association of Official Analytical Chemists (AOAC)**. Total carbohydrate content was determined by difference, and the energy value was calculated using the Atwater factors:

Energy (kcal/100g) = (4 x Protein) + (4 x Carbohydrate) + (9 x Lipid)

### 3.4 Phytochemical and Anti-nutrient Analysis

The anti-nutritional factors were quantified using spectrophotometric methods at the **Department of Biochemistry, University of Uyo**. Genesys 50, manufactured by Thermo Fisher Scientific, was used for the spectrophotometric analysis.

#### Total Cyanide

Determined using the alkaline picrate method.

#### Linamarin and Amygdalin

Quantified via specific enzymatic hydrolysis followed by spectrophotometric measurement of released cyanide.

#### Phytate and Oxalate

Measured using the methods described by Lucas and Markakas (1975) and Dye (1956), respectively.

##### Linamarase: UV spectrophotometric analysis

1g of sample was diluted with 50ml of 0.1M phosphate buffer at pH 6.0, kept for 5 min and centrifuged at 5000 rpm for 15 min. Linamarase was precipitated by 60% ammonium sulphate. The activity of the linamarase was estimated using 0.2ml of 0.2μmol linamarin, 1ml of enzyme and 0.8ml of phosphate buffer adjusted to pH 6.0. The mixtures were incubated for 15 mins at 35°C. The reaction was stopped by addition of 1ml of 0.1M NaOH and then neutralized with 1ml of 0.1M HCl. 1ml of 1% chloramine-T was added followed by 3ml of pyridine barbituric acid reagent. The mixture was read at 500nm (Nambisan and Sundaresan, 1994).

A standard calibration curve was prepared using known concentrations of the substrate (0 to 2.5 mmol/ml). The resulting linear regression equation was y = 0.0226x + 0.0004, with a coefficient of determination (R^2^) of 0.9915, ensuring high analytical precision.

### 3.5 Determination of Vitamins and Antioxidants

#### Vitamins C and E

Vitamin C was determined using the 2,6-dichlorophenolindophenol (DCPIP) titrimetric method, while Vitamin E was quantified spectrophotometrically using the Emmerie-Engel reaction.

#### Total Phenolic Content (TPC)

Assayed using the **Folin-Ciocalteu reagent** method, with results expressed as mg Gallic Acid Equivalents (GAE) per 100g.

#### Total Flavonoid Content (TFC)

Determined via the aluminum chloride colorimetric assay, with results expressed as mg Quercetin Equivalents (QE) per 100g.

### 3.6 Statistical Analysis

Data were expressed as Mean +/- Standard Deviation (SD) of triplicate determinations. Statistical significance was analyzed using **One-Way Analysis of Variance (ANOVA)** followed by **Tukey’s post-hoc test** for multiple comparisons. A p-value of < 0.05 was considered statistically significant. All analyses were performed using SPSS version 26.0

### 3.7 Ethical Approval

Ethical approval was given by the Faculty of Basic Medical Sciences, University of Uyo with the ethical number: UU FBMSREC 2025 020.

## 4. Results

All biochemical and nutritional parameters measured across the five processing stages (Fresh, Boiled, D1 (24h), D2 (48h), and D3 (72h)) are expressed as mg/100g of fresh weight (FW), unless otherwise stated.

The study first examines how boiling and varying durations of fermentation (24, 48, and 72 hours) alter the proximate composition, anti-nutrient levels, vitamin content, and antioxidant properties of cassava tubers compared to the fresh state. Statistical analyses (one-way ANOVA followed by Tukey HSD post-hoc tests) were then applied to confirm significant changes across processing stages.

### 4.1 Analysis of cassava composition data

#### 4.1.1 Proximate Composition

**Table 1:** Average Proximate Composition of Cassava Tuber Across Processing Stages (Values are mean ± SD, n = 3; g/100g except Energy in kcal/mol. Means in the same row with different superscript letters are significantly different (one-way ANOVA followed by Tukey HSD post-hoc test, p < 0.05).)

| Parameter | Fresh | Boiled | D1 | D2 | D3 |
| --- | --- | --- | --- | --- | --- |
| Moisture | 22.86 ± 0.09<sup>a</sup> | 20.12 ± 0.09<sup>d</sup> | 20.70 ± 0.06<sup>cd</sup> | 21.81 ± 0.10<sup>b</sup> | 21.68 ± 0.48<sup>b</sup> |
| Protein | 1.08 ± 0.08<sup>ab</sup> | 0.87 ± 0.06<sup>b</sup> | 0.98 ± 0.12<sup>ab</sup> | 1.18 ± 0.23<sup>a</sup> | 1.28 ± 0.27<sup>a</sup> |
| Lipid | 0.34 ± 0.06<sup>a</sup> | 0.14 ± 0.02<sup>d</sup> | 0.63 ± 0.03<sup>b</sup> | 0.57 ± 0.04<sup>bc</sup> | 0.48 ± 0.09<sup>c</sup> |
| Fiber | 4.13 ± 0.14<sup>ab</sup> | 4.12 ± 0.10<sup>b</sup> | 4.42 ± 0.11<sup>a</sup> | 4.29 ± 0.10<sup>ab</sup> | 4.01 ± 0.12<sup>b</sup> |
| Ash | 1.13 ± 0.07<sup>a</sup> | 0.78 ± 0.03<sup>d</sup> | 1.40 ± 0.02<sup>a</sup> | 1.36 ± 0.06<sup>a</sup> | 1.30 ± 0.02<sup>a</sup> |
| Carbohydrate | 70.47 ± 0.12<sup>d</sup> | 73.98 ± 0.18<sup>a</sup> | 71.86 ± 0.35<sup>bc</sup> | 70.79 ± 0.21<sup>cd</sup> | 71.26 ± 0.35<sup>bc</sup> |
| Energy (kcal/mol) | 289.38 ± 0.15<sup>d</sup> | 300.68 ± 0.80<sup>a</sup> | 296.72 ± 0.42<sup>b</sup> | 292.97 ± 0.30<sup>c</sup> | 294.47 ± 1.13<sup>bc</sup> |

*Superscript letters (<sup>a</sup>, <sup>b</sup>, <sup>c</sup>, <sup>d</sup>, etc.) indicate statistically significant differences among processing stages within each parameter row (Tukey HSD post-hoc test, $p < 0.05$ ).*
*Means sharing the same letter are not significantly different from each other.*
*D1: Boiled and Fermented for 24hrs*
*D2: Boiled and Fermented for 48hrs*
*D3: Boiled and Fermented for 72hrs*
**Note: All concentrations are calculated on a fresh weight (FW) basis. (Units are expressed as mg/100g fresh weight).**
*D1: Boiled and Fermented for 24hrs; D2: Boiled and Fermented for 48hrs*

## 2. Analysis of cassava composition data

### 2.1 Proximate Composition

As presented in Table 1, one-way ANOVA confirmed significant differences across the five processing stages (Fresh, Boiled, D1, D2, D3) for all proximate parameters (p < 0.05, partial η² = 0.78–0.97), indicating that processing substantially altered the nutritional composition. Post-hoc comparisons showed that boiling significantly reduced moisture content compared to fresh cassava (p < 0.001), with fermented stages (especially D2 and D3) showing partial recovery but still significantly lower than fresh (p < 0.05). Carbohydrate and energy content increased significantly after boiling due to moisture loss (concentration effect; p < 0.001 vs. fresh), then decreased during fermentation toward fresh values. Lipid, ash, and protein also changed significantly, with boiled cassava having the lowest lipid and ash levels, while fermented stages generally returned closer to fresh values. Fiber remained relatively stable across stages, with only minor significant differences. These results demonstrate that boiling and fermentation induce real, statistically meaningful changes in proximate composition rather than random variation.

Descriptively, as presented in Figure 1, the moisture content was highest in fresh cassava tuber (22.86 g/100g). It decreased after boiling (20.12 g/100g) and stayed relatively low in the fermented samples (D1: 20.7 g/100g, D2: 21.81 g/100g, D3: 21.68 g/100g). This reduction in moisture is expected after boiling and during fermentation due to water loss and changes in the tuber structure.

**Figure 1:**
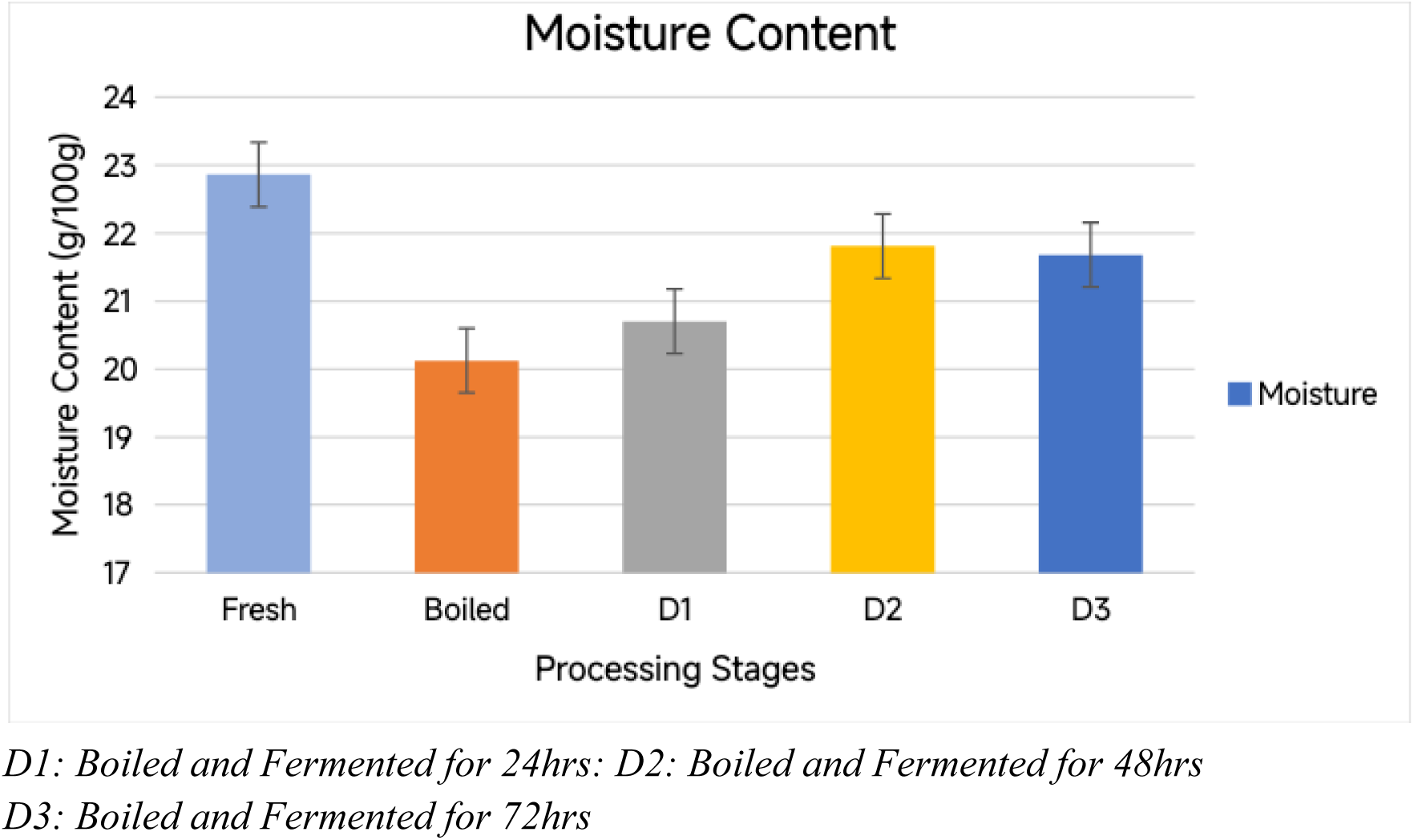
Moisture Content Across Processing Stages of Cassava Tuber. Note: All concentrations are calculated on a fresh weight (FW) basis. (Units are expressed as mg/100g fresh weight).

As shown in Figure 2 above, fiber content remained fairly stable across all processing stages, ranging from 4.01 to 4.42 g/100g. It was slightly higher in the boiled + 24hr fermented sample (D1) compared to the fresh and boiled cassava tuber, then gradually decreased in D2 and D3. Overall, processing had only a minor effect on fiber content.

**Figure 2:**
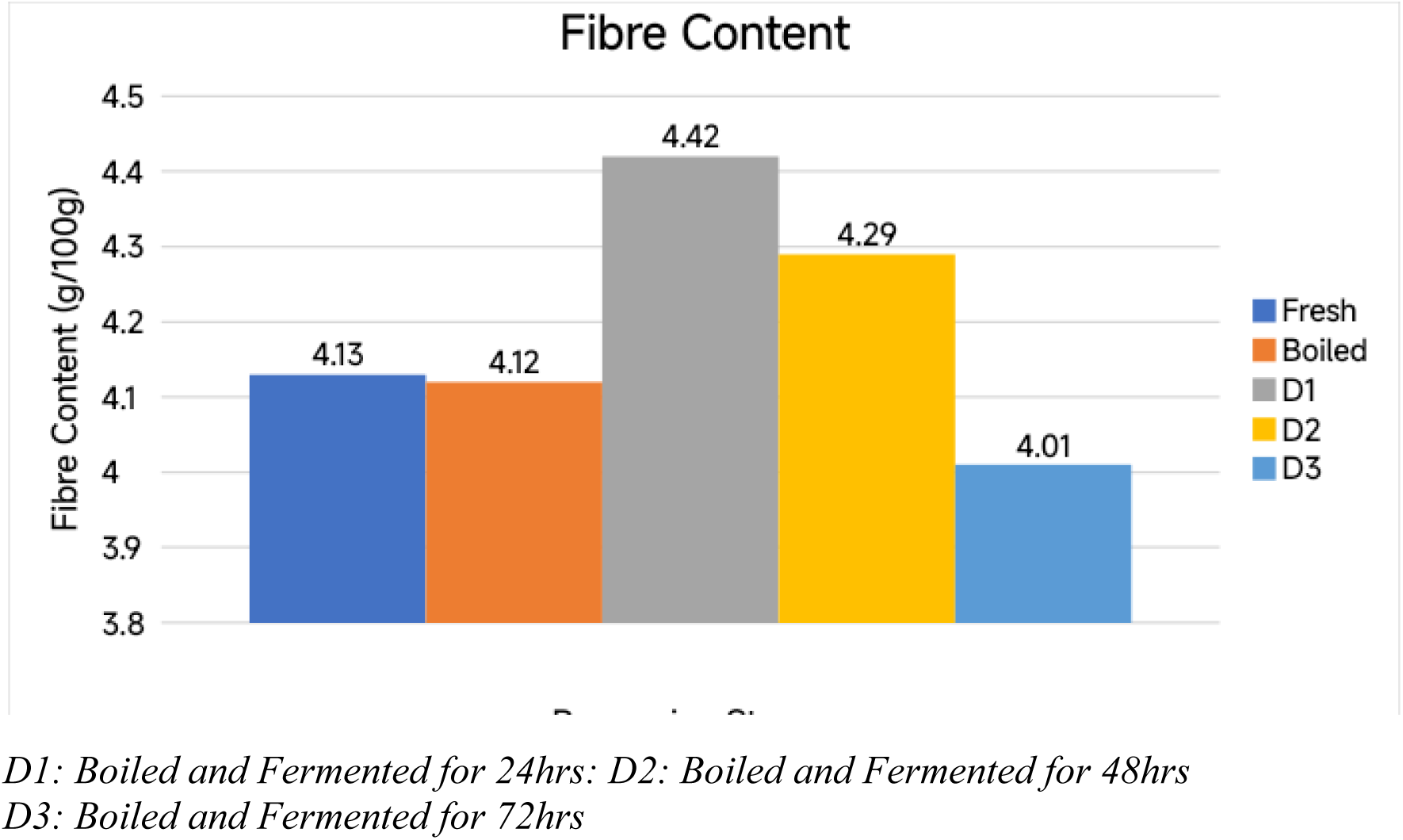
Fiber Content Across Processing Stages of Cassava Tuber. Note: All concentrations are calculated on a fresh weight (FW) basis. (Units are expressed as mg/100g fresh weight).

As shown in Figure 3 above, carbohydrate content was lowest in fresh cassava tuber (70.47 g/100g). It increased clearly after boiling (73.98 g/100g), likely due to moisture loss, concentrating the carbohydrates. The fermented samples (D1, D2, D3) showed values closer to fresh cassava tuber, suggesting that fermentation partially reversed this concentration effect.

**Figure 3:**
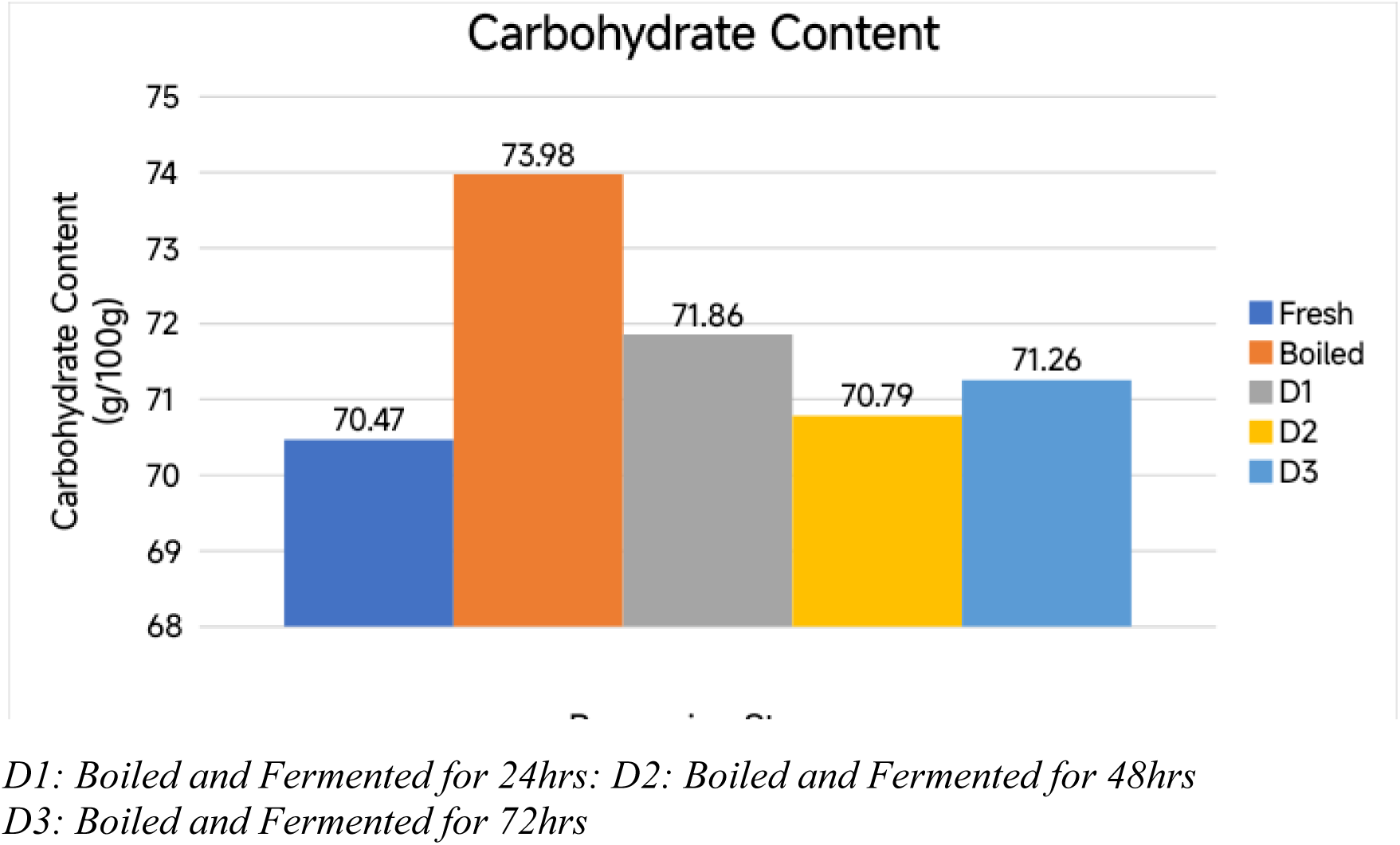
Carbohydrate Content Across Processing Stages of Cassava Tuber. Note: All concentrations are calculated on a fresh weight (FW) basis. (Units are expressed as mg/100g fresh weight).

As shown in Figure 4 above, the energy value followed a similar pattern to that of carbohydrate. It was lowest in fresh cassava tuber (289.38 kcal/mol), peaked after boiling (300.68 kcal/mol), and then decreased slightly in the fermented samples (292.97–296.72 kcal/mol). The higher energy in boiled cassava tuber is mainly due to the increased proportion of carbohydrates after moisture is reduced.

**Figure 4:**
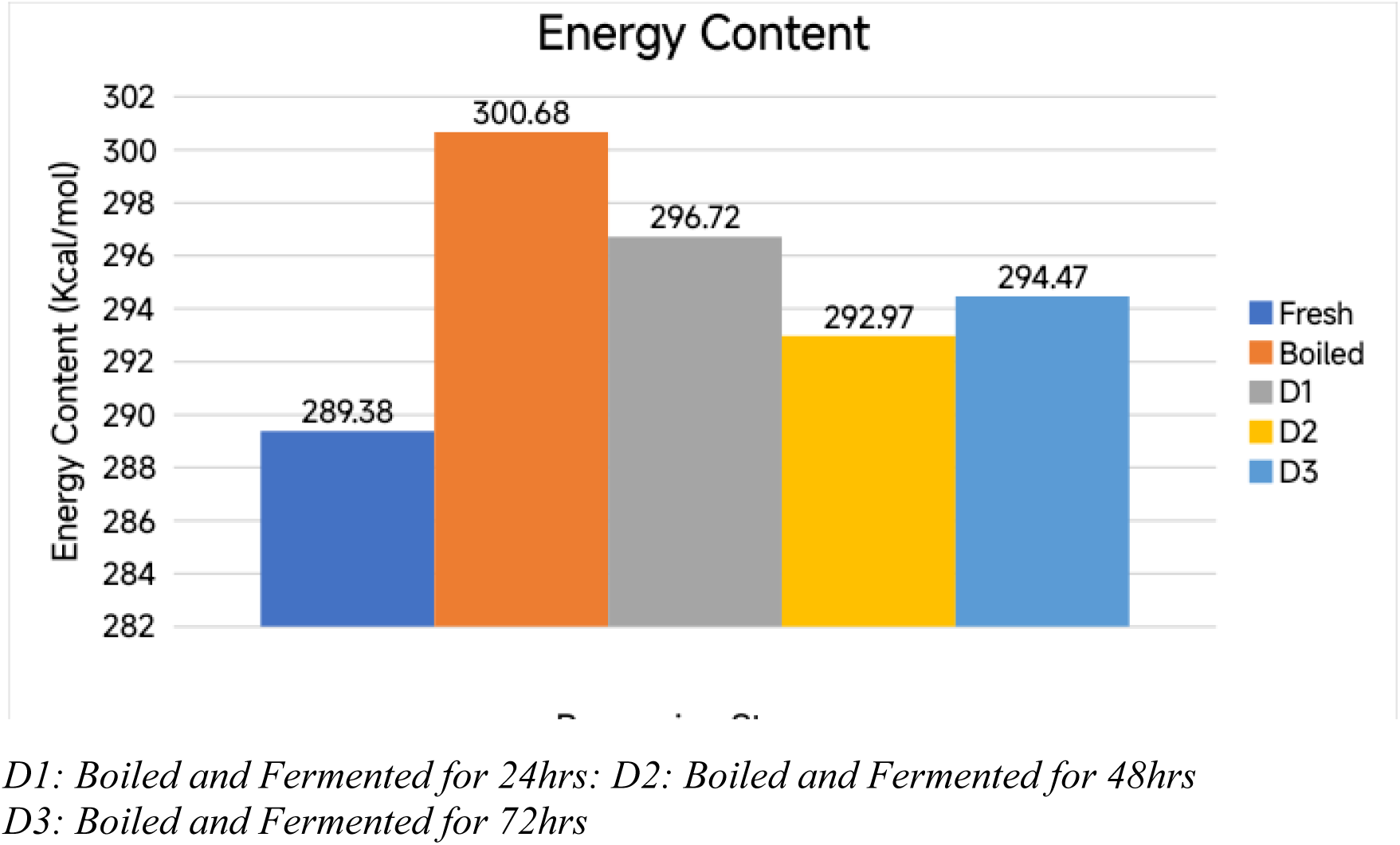
Energy Content Across Processing Stages of Cassava Tuber. Note: All concentrations are calculated on a fresh weight (FW) basis. (Units are expressed as mg/100g fresh weight)

From Figure 5, protein, lipid, and ash showed small variations across the processing stages. Protein was lowest in boiled cassava tuber (0.87 g/100g) and gradually increased in the fermented samples (up to 1.28 g/100g in D3). Lipid content was lowest after boiling (0.14 g/100g) and increased noticeably in the fermented samples (0.48–0.63 g/100g). Ash content dropped after boiling (0.78 g/100g) and returned to levels similar to those of fresh cassava tuber in the fermented samples. These small changes indicate that fermentation has a mild positive effect on protein and lipid content.

**Figure 5:**
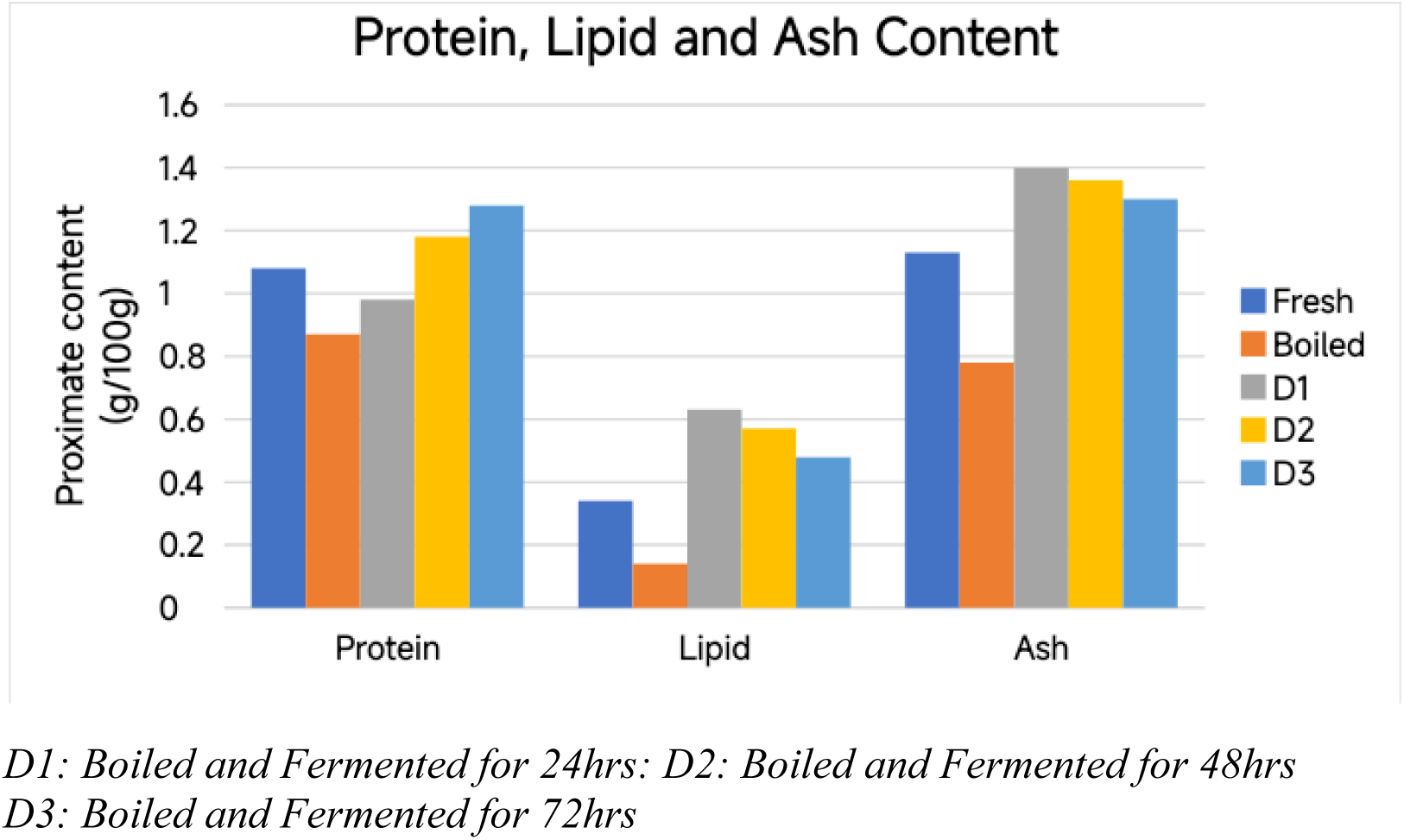
Protein, Lipid, and Ash Content Across Processing Stages of Cassava Tuber. Note: All concentrations are calculated on a fresh weight (FW) basis. (Units are expressed as mg/100g fresh weight)

### 2.2 Anti-nutrient Composition

As presented in Table 2, one-way ANOVA revealed highly significant differences across the five processing stages (Fresh, Boiled, D1, D2, D3) for all anti-nutrient parameters (total cyanide, oxalate, phytate, amygdalin, and linamarin) (p < 0.001, partial η² ≈ 1.0), confirming that processing induces dramatic and reliable reductions in these compounds. Post-hoc comparisons (Tukey HSD) showed that every stage was significantly different from the others (p < 0.001 for all pairwise tests), with no overlap between groups. Total cyanide decreased progressively from 98.15 mg/100g in fresh cassava to 59.02 mg/100g after boiling, then sharply to 3.67 mg/100g in D1, 1.70 mg/100g in D2, and just 0.54 mg/100g in D3. Similar stepwise and highly significant reductions were observed for oxalate, phytate, amygdalin, and linamarin, with the longest fermentation (D3) consistently yielding the lowest levels. These results demonstrate that boiling removes a substantial portion of anti-nutrients, while fermentation further detoxifies the cassava in a time-dependent manner, with each additional fermentation stage producing statistically meaningful improvement.

**Table 2:** Average Anti-nutrient Composition of Cassava Tuber Across Processing Stages (Values are mean ± SD, n = 3; mg/100g. Means in the same row with different superscript letters are significantly different (one-way ANOVA followed by Tukey HSD post-hoc test, p < 0.05).)

| Parameter | Fresh | Boiled | D1 | D2 | D3 |
| --- | --- | --- | --- | --- | --- |
| Total Cyanide | 98.15 ± 0.05<sup>a</sup> | 59.02 ± 0.64<sup>b</sup> | 3.67 ± 0.06<sup>c</sup> | 1.70 ± 0.12<sup>d</sup> | 0.54 ± 0.05<sup>e</sup> |
| Oxalate | 44.26 ± 0.06<sup>a</sup> | 16.81 ± 0.07<sup>b</sup> | 6.18 ± 0.17<sup>c</sup> | 4.21 ± 0.13<sup>d</sup> | 2.13 ± 0.05<sup>e</sup> |
| Phytate | 304.24 ± 0.04<sup>a</sup> | 135.21 ± 0.27<sup>b</sup> | 18.68 ± 0.06<sup>c</sup> | 11.24 ± 0.14<sup>d</sup> | 7.14 ± 0.06<sup>e</sup> |
| Amygdalin | 1.86 ± 0.02<sup>a</sup> | 0.54 ± 0.01<sup>b</sup> | 0.41 ± 0.01<sup>c</sup> | 0.32 ± 0.01<sup>d</sup> | 0.26 ± 0.01<sup>e</sup> |
| Linamarin | 136.44 ± 0.05<sup>a</sup> | 11.10 ± 0.18<sup>b</sup> | 5.16 ± 0.08<sup>c</sup> | 3.72 ± 0.06<sup>d</sup> | 1.15 ± 0.02<sup>e</sup> |

*Superscript letters (<sup>a</sup>, <sup>b</sup>, <sup>c</sup>, <sup>d</sup>, <sup>e</sup>) indicate statistically significant differences among processing stages within each parameter row (Tukey HSD post-hoc test, $p < 0.05$ ).*
*Means sharing the same letter are not significantly different from each other.*
*D1: Boiled and Fermented for 24hrs*
*D2: Boiled and Fermented for 48hrs*
*D3: Boiled and Fermented for 72hrs*
*D1: Boiled and Fermented for 24hrs; D2: Boiled and Fermented for 48hrs*

**Figure 6:**
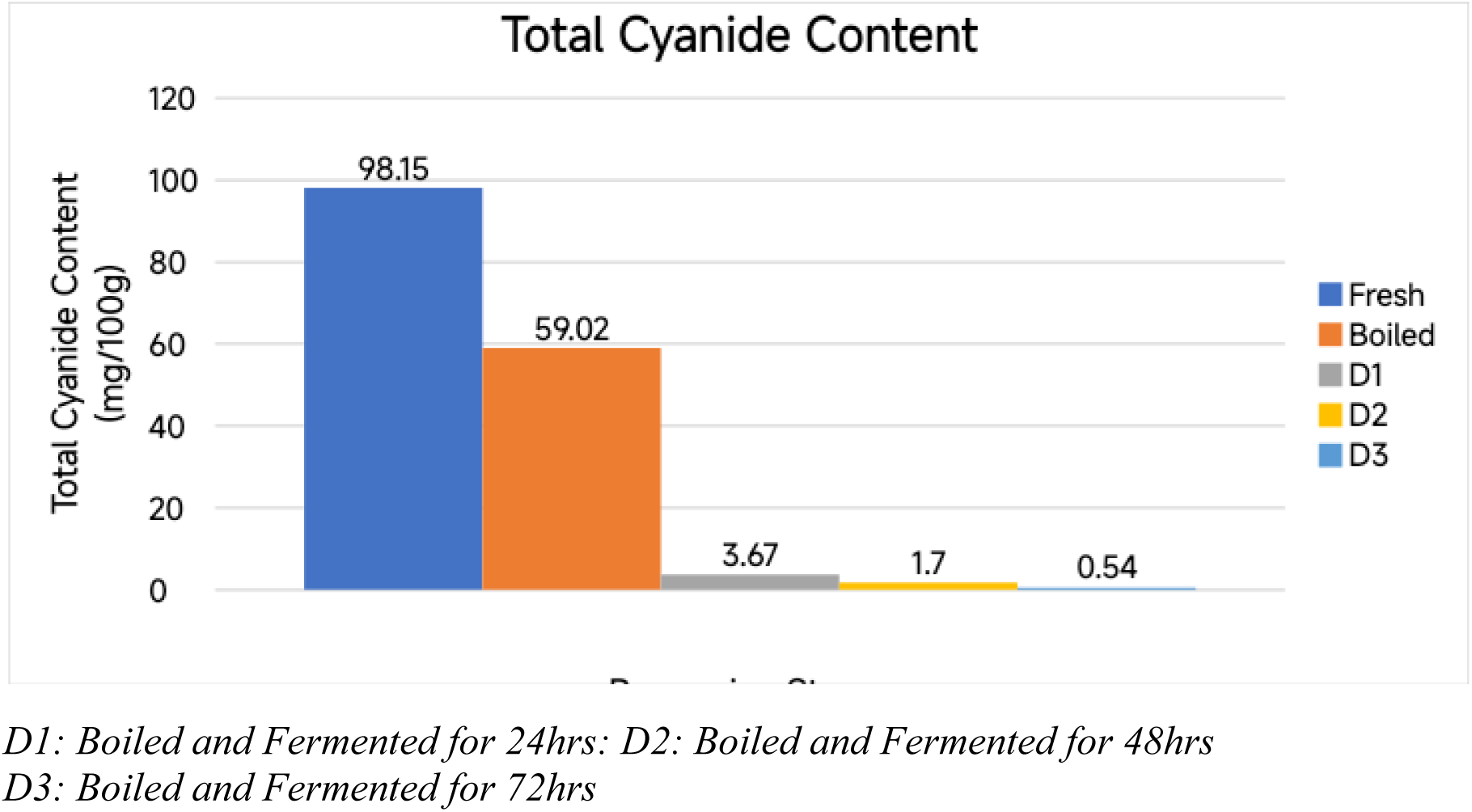
Total Cyanide Content Across Processing Stages of Cassava Tuber. Note: All concentrations are calculated on a fresh weight (FW) basis. (Units are expressed as mg/100g fresh weight).

Descriptively, as presented in Figure 6, total cyanide was very high in fresh cassava tuber (98.15 mg/100g). It decreased after boiling to 59.02 mg/100g, and then dropped dramatically during fermentation, reaching 3.67 mg/100g in D1, 1.7 mg/100g in D2, and only 0.54 mg/100g in D3. This shows that fermentation, especially longer fermentation, is highly effective at removing cyanide from cassava tuber.

**Figure 7:**
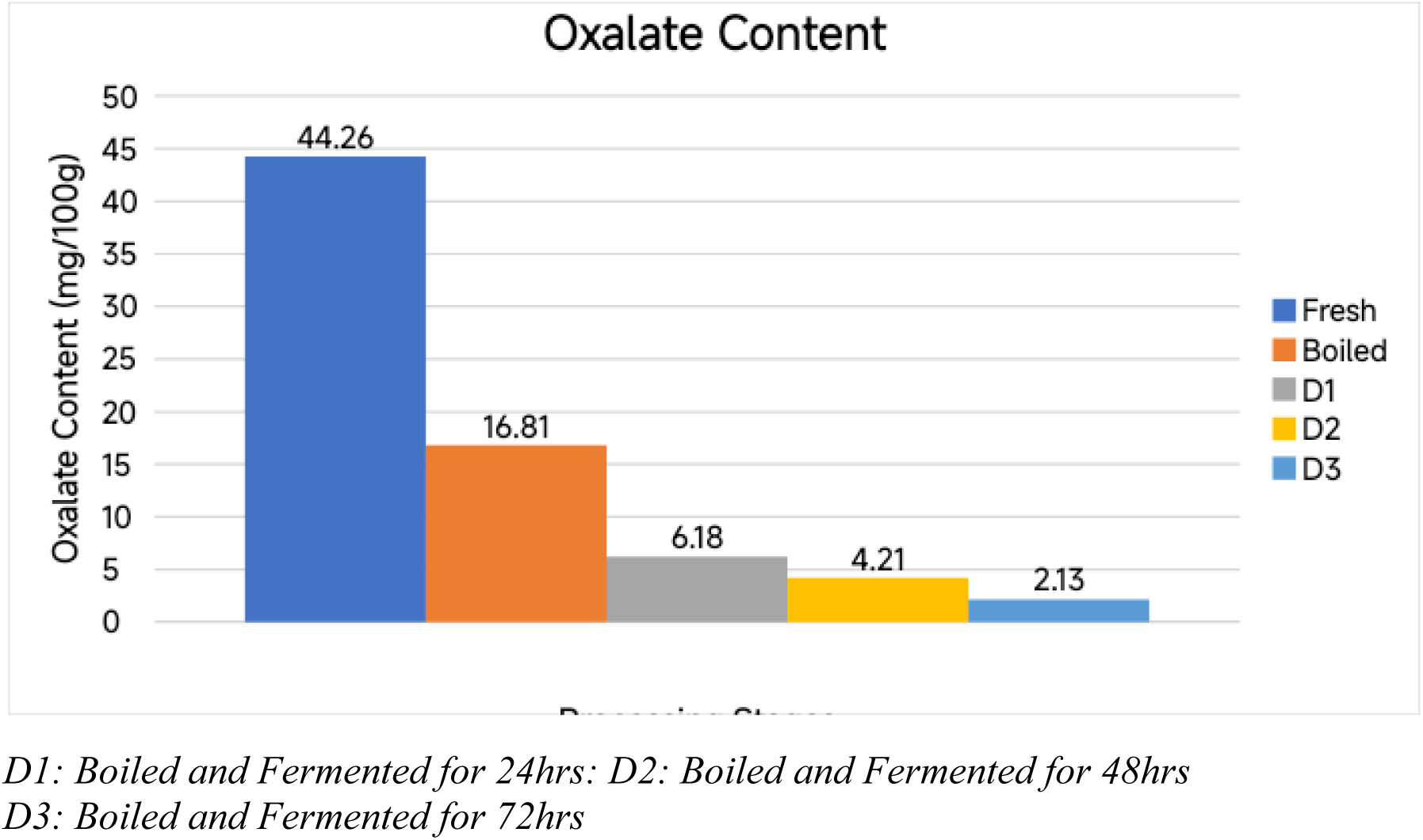
Oxalate Content Across Processing Stages of Cassava Tuber. Note: All concentrations are calculated on a fresh weight (FW) basis. (Units are expressed as mg/100g fresh weight)

Oxalate content started at 44.26 mg/100g in fresh cassava tuber and decreased to 16.81 mg/100g after boiling. It continued to fall steadily during fermentation (6.18 → 4.21 → 2.13 mg/100g in D1, D2, and D3). The consistent downward trend shows that both boiling and fermentation help reduce oxalate levels.

**Figure 8:**
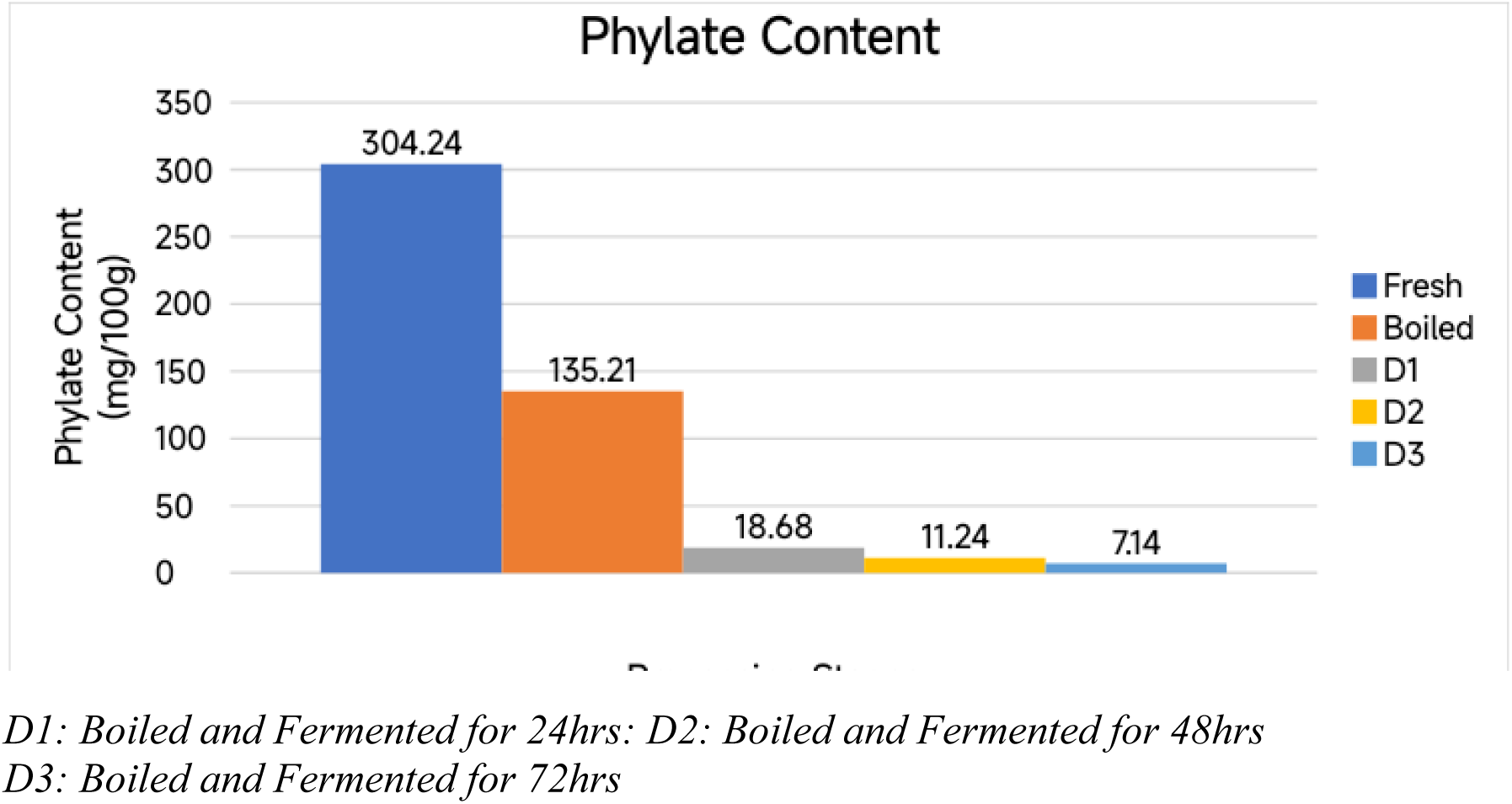
Phytate Content Across Processing Stages of Cassava Tuber. Note: All concentrations are calculated on a fresh weight (FW) basis. (Units are expressed as mg/100g fresh weight)

Phytate was extremely high in fresh cassava tuber (304.24 mg/100g). Boiling reduced it to 135.21 mg/100g, and fermentation caused a very sharp further decrease (18.68 → 11.24 → 7.14 mg/100g in D1, D2, D3). This indicates that fermentation is particularly effective at breaking down phytate in cassava.

**Figure 9:**
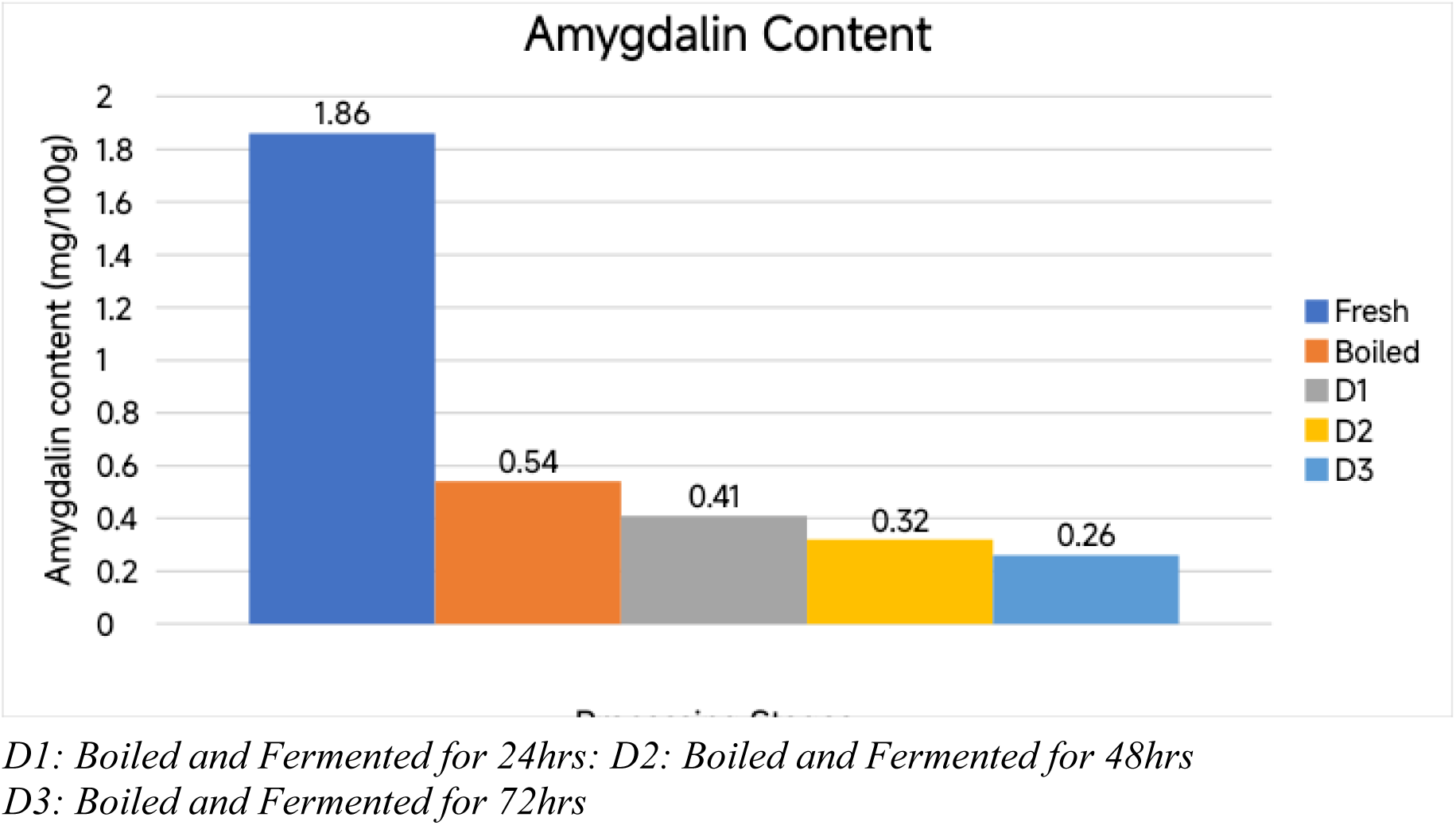
Amygdalin Content Across Processing Stages of Cassava Tuber. Note: All concentrations are calculated on a fresh weight (FW) basis. (Units are expressed as mg/100g fresh weight)

Amygdalin levels were low even in fresh cassava tuber (1.86 mg/100g). They decreased further after boiling (0.54 mg/100g) and continued to decline gradually during fermentation (0.41 → 0.32 → 0.26 mg/100g). Although the starting amount was small, processing still reduced amygdalin consistently.

**Figure 10:**
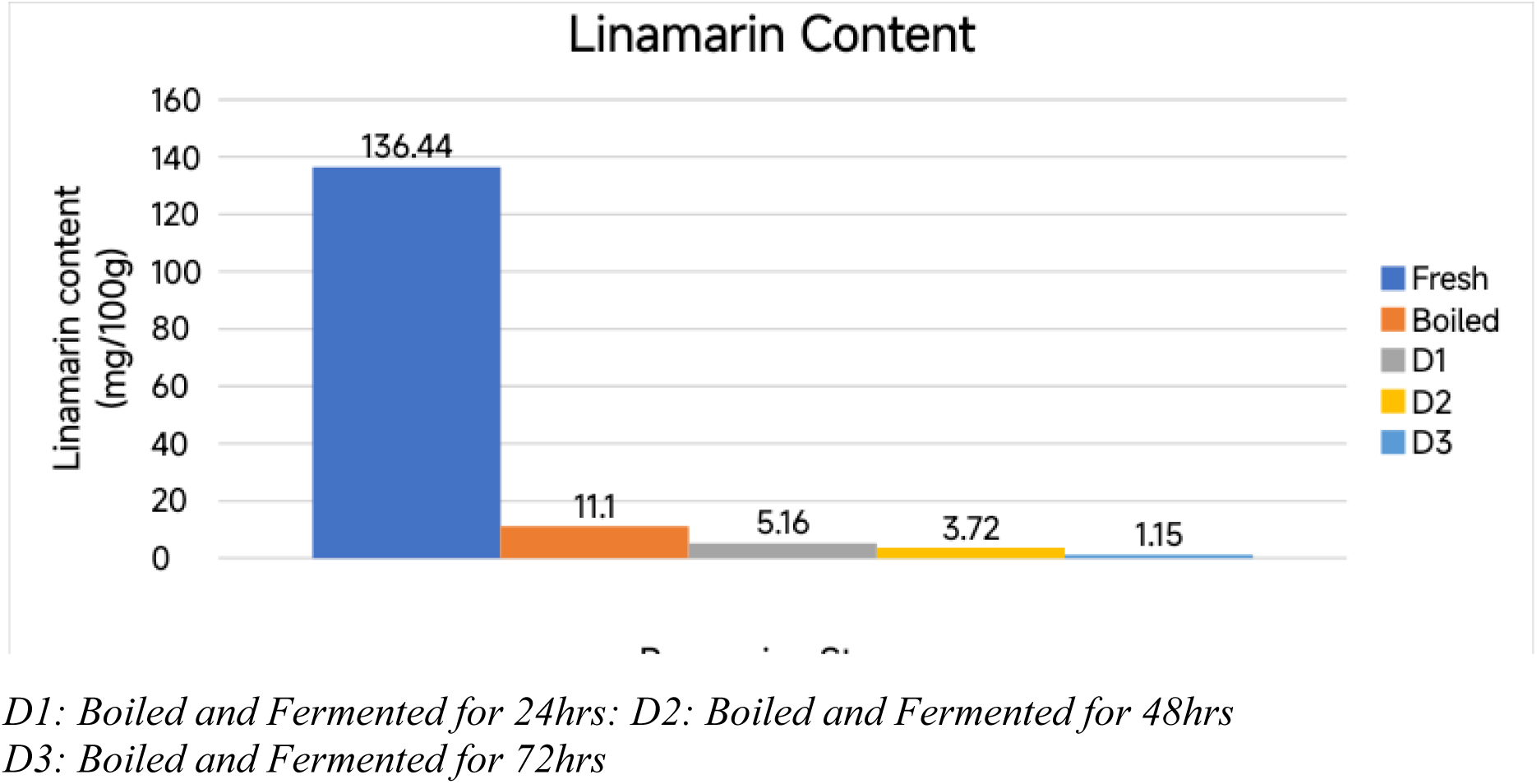
Linamarin Content Across Processing Stages of Cassava Tuber. Note: All concentrations are calculated on a fresh weight (FW) basis. (Units are expressed as mg/100g fresh weight)

Linamarin was high in fresh cassava tuber (136.44 mg/100g) and dropped sharply after boiling (11.1 mg/100g). Fermentation caused additional reduction (5.16 → 3.72 → 1.15 mg/100g in D1, D2, D3). Like total cyanide, linamarin showed a very strong decrease with boiling and especially with fermentation.

### 2.3 Vitamin Composition

**Table 3:** Average Vitamin Composition of Cassava Tuber Across Processing Stages (Values are mean ± SD, n = 3; mg/100g. Means in the same row with different superscript letters are significantly different (one-way ANOVA followed by Tukey HSD post-hoc test, p < 0.05).

| Parameter | Fresh | Boiled | D1 | D2 | D3 |
| --- | --- | --- | --- | --- | --- |
| Vitamin C | 26.71 ± 0.01<sup>a</sup> | 1.82 ± 0.02<sup>e</sup> | 12.33 ± 0.01<sup>b</sup> | 10.16 ± 0.02<sup>c</sup> | 7.64 ± 0.03<sup>d</sup> |
| Vitamin E | 0.29 ± 0.01<sup>ab</sup> | 0.81 ± 0.02<sup>a</sup> | 0.22 ± 0.02<sup>bc</sup> | 0.17 ± 0.04<sup>bc</sup> | 0.11 ± 0.01<sup>c</sup> |

*Superscript letters (<sup>a</sup>, <sup>b</sup>, <sup>c</sup>, <sup>d</sup>, <sup>e</sup>) indicate statistically significant differences among processing stages within each parameter row (Tukey HSD post-hoc test, $p < 0.05$ ).*
*Means sharing the same letter are not significantly different from each other*
*D1: Boiled and Fermented for 24hrs*
*D2: Boiled and Fermented for 48hrs*
*D3: Boiled and Fermented for 72hrs*
***Note: All concentrations are calculated on a fresh weight (FW) basis. (Units are expressed as mg/100g fresh weight)***
*D1: Boiled and Fermented for 24hrs; D2: Boiled and Fermented for 48hrs*

One-way ANOVA revealed highly significant differences across the five processing stages for vitamin C (p < 0.001, partial η² ≈ 1.0) and a significant difference for vitamin E (p < 0.05, partial η² = 0.587). Post-hoc comparisons (Tukey HSD) confirmed that all stages were significantly different from each other for vitamin C (p < 0.001 for all pairwise comparisons), with fresh cassava tuber showing the highest content (26.71 mg/100g), a dramatic reduction after boiling (1.82 mg/100g), and progressive partial recovery during fermentation (12.33 mg/100g in D1, 10.16 mg/100g in D2, and 7.64 mg/100g in D3). For vitamin E, levels remained very low overall; boiling caused a significant increase (0.81 mg/100g) compared to fresh (0.29 mg/100g), likely due to concentration from moisture loss, while fermentation resulted in a gradual decline (0.22 mg/100g in D1 to 0.11 mg/100g in D3), with D3 significantly lower than boiled (p < 0.05). These findings indicate that boiling destroys most of the vitamin C present in fresh cassava tuber, but fermentation allows partial restoration in a time-dependent manner, whereas vitamin E exhibits only minor and inconsistent changes across processing stages.

Descriptively, as presented in Figure 11, fresh cassava tuber contained a substantial amount of vitamin C (26.71 mg/100g). After boiling, the level fell sharply to only 1.82 mg/100g, showing that boiling destroys most of the vitamin C. During fermentation, vitamin C increased again (12.33 mg/100g in D1, 10.16 mg/100g in D2, and 7.64 mg/100g in D3), but it never returned to the fresh level. This pattern indicates that while fermentation helps restore some vitamin C, significant loss still occurs during the overall processing.

**Figure 11:**
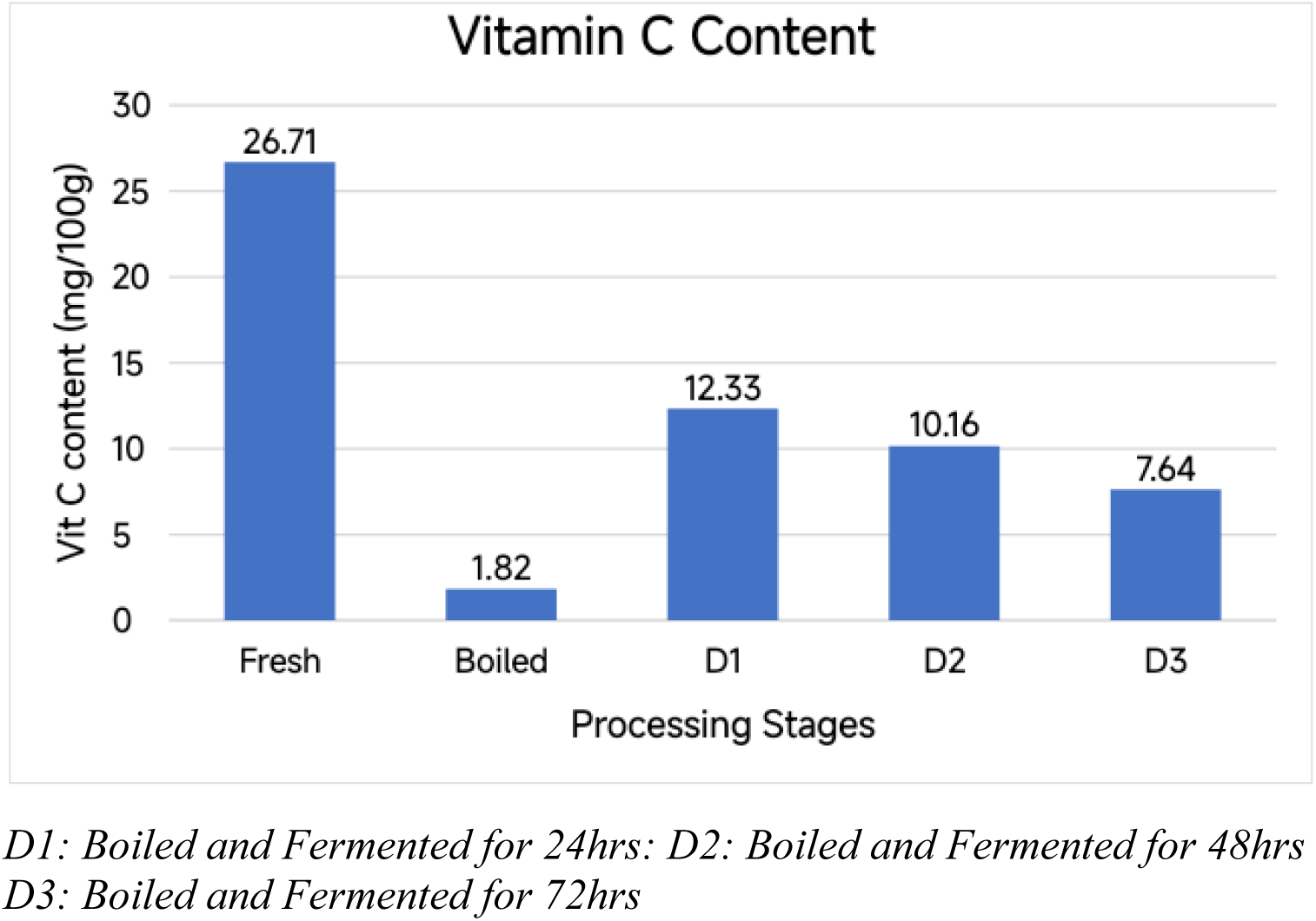
Vitamin C Content Across Processing Stages. Note: All concentrations are calculated on a fresh weight (FW) basis. (Units are expressed as mg/100g fresh weight)

**Figure 12:**
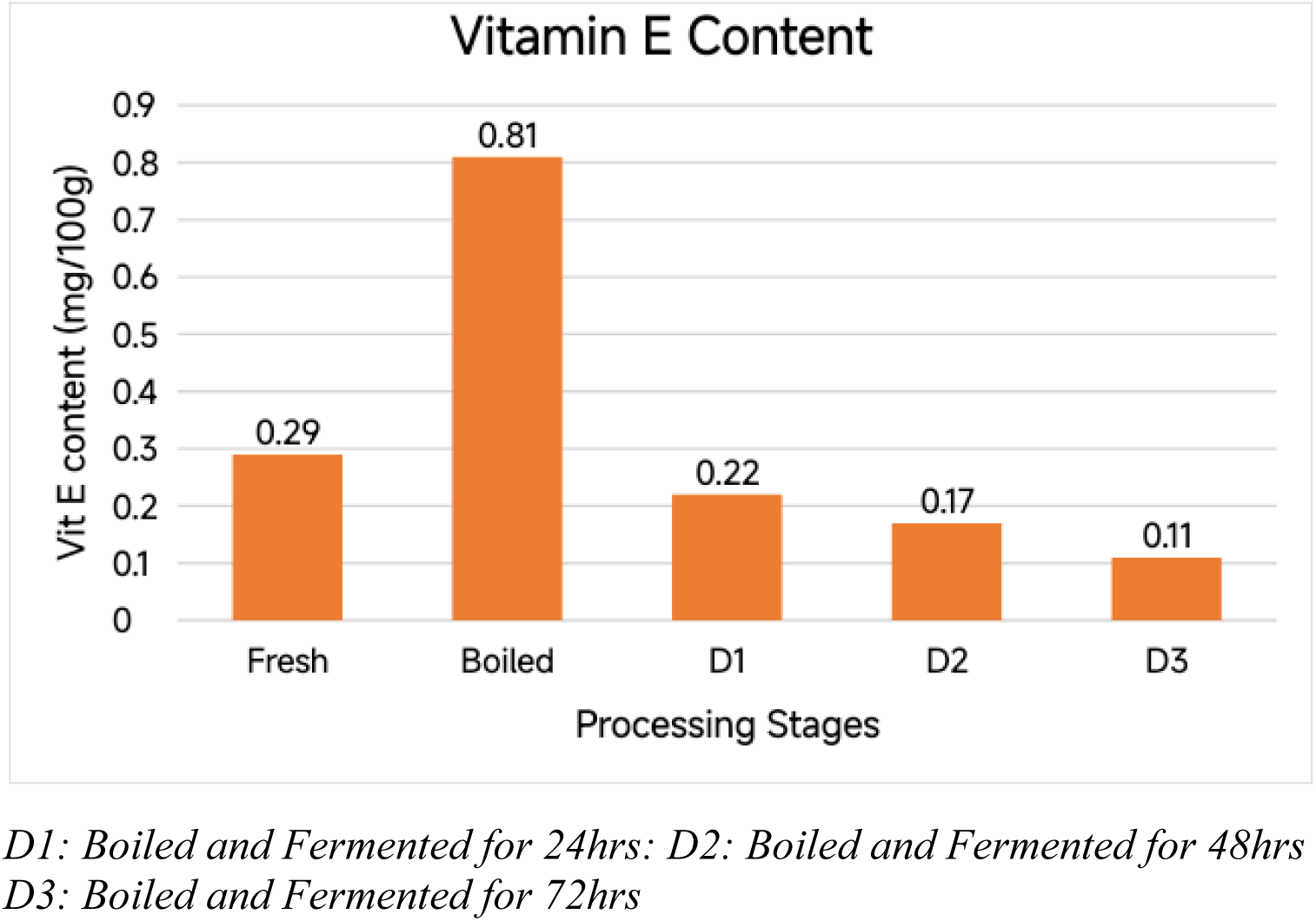
Vitamin E Content Across Processing Stages of Cassava Tuber. Note: All concentrations are calculated on a fresh weight (FW) basis. (Units are expressed as mg/100g fresh weight)

Vitamin E was present in very low amounts in fresh cassava tuber (0.29 mg/100g). It increased slightly after boiling (0.81 mg/100g), possibly due to concentration from moisture loss. In the fermented samples, it then decreased steadily (0.22 → 0.17 → 0.11 mg/100g in D1, D2, and D3). Overall, vitamin E levels remained very low throughout processing and showed only minor changes.

### 2.4 Antioxidant Composition

**Table 4:**
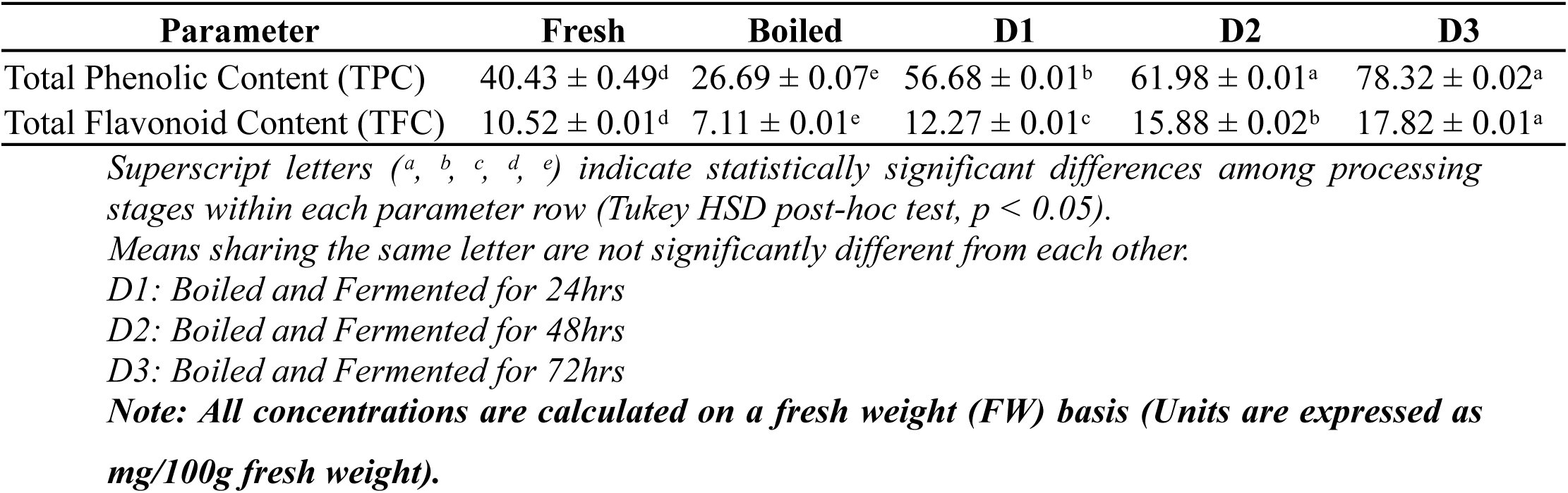
Average Antioxidant Composition of Cassava Tuber Across Processing Stages (Values are mean ± SD, n = 3; TPC in mg GAE/100g, TFC in mg QE/100g. Means in the same row with different superscript letters are significantly different (one-way ANOVA followed by Tukey HSD post-hoc test, p < 0.05).

One-way ANOVA revealed highly significant differences across the five processing stages for both total phenolic content (TPC) and total flavonoid content (TFC) (p < 0.001, partial η² ≈ 1.0). Post-hoc comparisons (Tukey HSD) confirmed that all stages were significantly different from each other for both parameters (p < 0.001 for most pairs). TPC decreased significantly after boiling (from 40.43 mg GAE/100g in fresh to 26.69 mg GAE/100g), likely due to heat-induced degradation, then increased progressively and substantially during fermentation (56.68 mg GAE/100g in D1, 61.98 mg GAE/100g in D2, and 78.32 mg GAE/100g in D3), with D2 and D3 showing the highest levels. TFC followed a very similar pattern: a significant drop after boiling (from 10.52 mg QE/100g in fresh to 7.11 mg QE/100g), followed by steady and significant increases during fermentation (12.27 mg QE/100g in D1, 15.88 mg QE/100g in D2, and 17.82 mg QE/100g in D3), with D3 being the highest. These results demonstrate that while boiling reduces antioxidant compounds, fermentation strongly enhances both total phenolics and flavonoids in a time-dependent manner, with longer fermentation (D3) producing the greatest antioxidant capacity.

**Figure 13:**
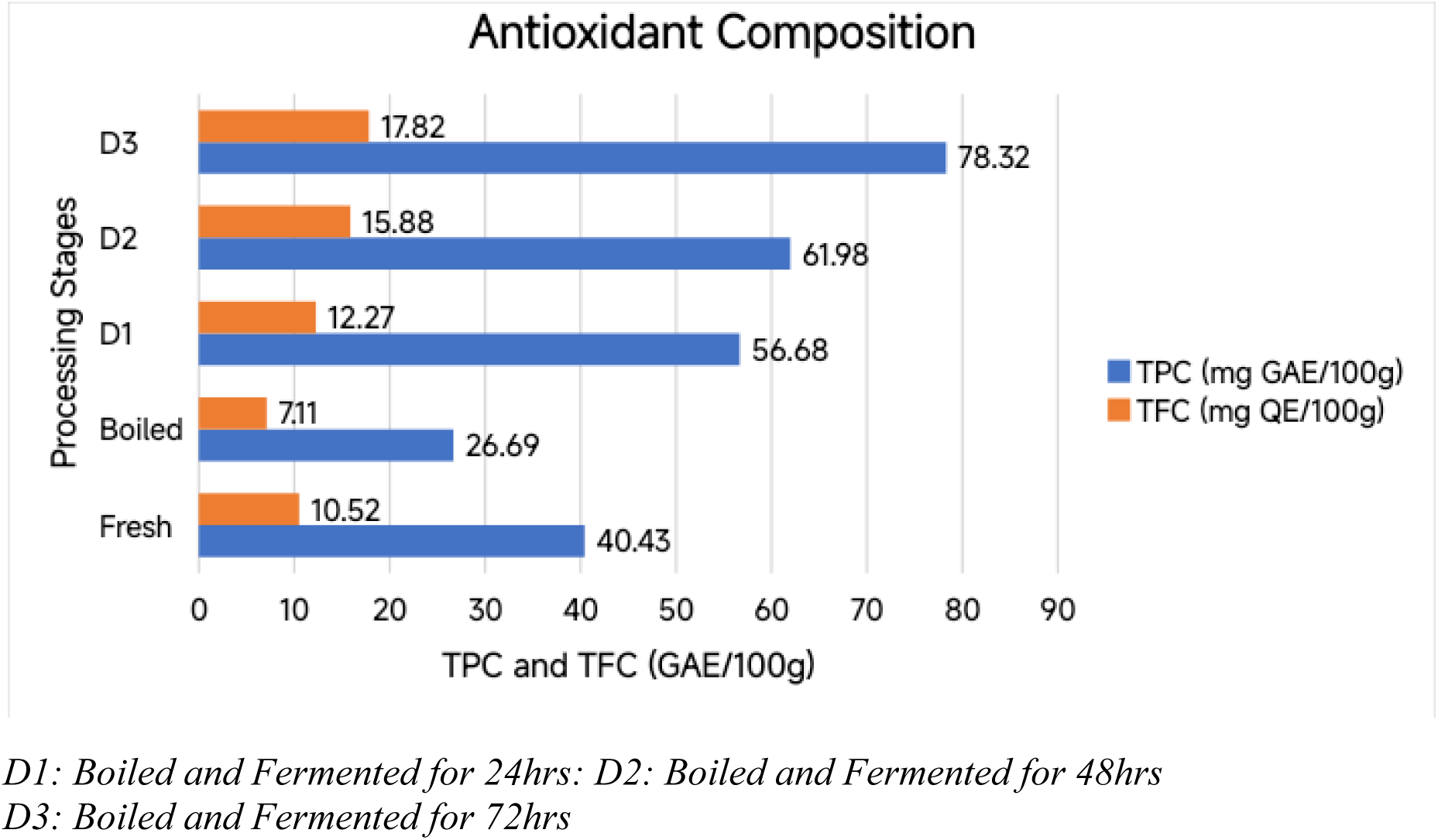
Total Phenolic Content (TPC) and Total Flavonoid Content (TFC) Across Processing Stages of Cassava Tuber (Units are expressed as mg/100g fresh weight) Note: All concentrations are calculated on a fresh weight (FW) basis.

Descriptively, total phenolic content (TPC) was 40.43 mg GAE/100g in fresh cassava tuber and dropped to 26.69 mg GAE/100g after boiling. During fermentation, TPC increased progressively and substantially (56.68 → 61.98 → 78.32 mg GAE/100g in D1, D2, and D3). Total flavonoid content (TFC) followed a very similar pattern: it decreased from 10.52 mg QE/100g in fresh cassava tuber to 7.11 mg QE/100g after boiling, then rose steadily during fermentation (12.27 → 15.88 → 17.82 mg QE/100g). These results show that while boiling reduces antioxidant compounds, fermentation strongly enhances both total phenolics and flavonoids, making fermented cassava tuber richer in antioxidant activity.

### 4.5 Linamarase Activity Across Sequential Processing Stages

Table 5 presents the enzymatic activity of linamarase across sequential processing stages. Activity was lowest in fresh tubers (1.6425 units/g) but increased significantly following the 10-minute boiling stage (9.4726 units/g), suggesting a temperature-induced release or activation of the enzyme. During the fermentation phase (D1–D3), activity remained elevated compared to the fresh state, peaking at 8.5168 units/g by 72 hours (D3). This sustained activity is critical for the near-complete hydrolysis of residual cyanogenic glycosides observed in the later stages of processing

**Table 5:** Linamarase Activity Across Sequential Processing Stages.

| Sample | Absorbance | Conc mmol/50 ml | Conc mmol/ml | Conc \mu mol/ml | Conc unit/min | Conc unit/g sample |
| --- | --- | --- | --- | --- | --- | --- |
| Fresh (F) | 0.0468 | 2.053 | 0.0411 | 41.06195 | 1.6425 | 1.6425 |
| Boiled (B) | 0.2680 | 11.841 | 0.2368 | 236.8142 | 9.4726 | 9.4726 |
| D1 (24h) | 0.2180 | 9.628 | 0.1926 | 192.5664 | 7.7027 | 7.7027 |
| D2 (48h) | 0.0722 | 3.177 | 0.0635 | 63.53982 | 2.5416 | 2.5416 |
| D3 (72h) | 0.2410 | 10.646 | 0.2129 | 212.9204 | 8.5168 | 8.5168 |

***Note:** Processing conditions included boiling (10 min, open pot) followed by sequential fermentation (D1=24h, D2=48h, D3=72h). One unit of linamarase activity is defined as the amount of enzyme required to release 1 μmol of HCN per minute under the specified assay conditions (pH 6.0, 35°C). Assay weight = 1g; Incubation time = 25 mins. Values are based on spectrophotometric analysis at 500nm.*

**Figure 14.**
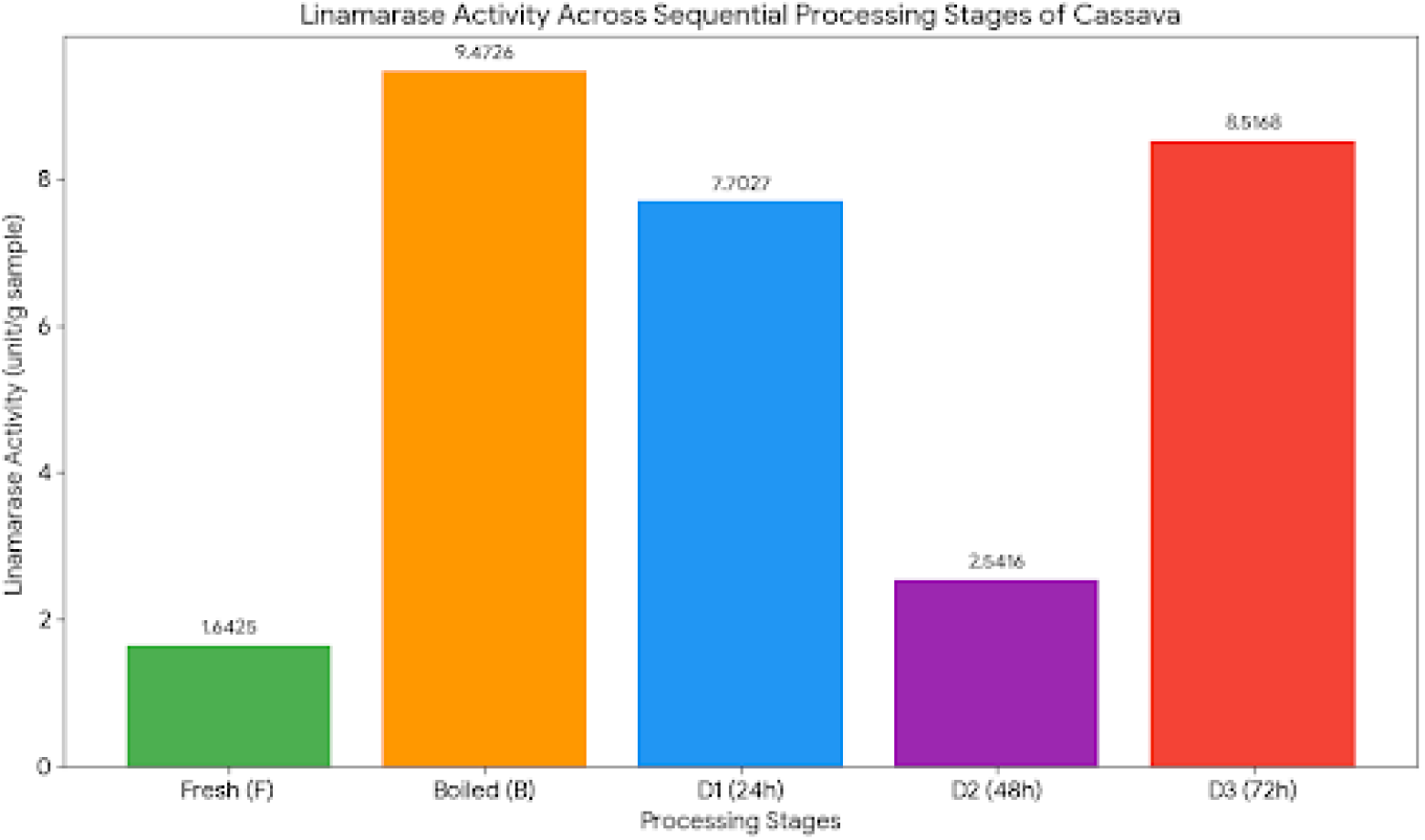
**above** illustrates the time-dependent resurgence of linamarase activity during fermentation. While boiling caused an initial spike in detectable activity, the subsequent 72-hour fermentation period (D3) maintained high enzymatic potency. This profile indicates that the fermentation environment facilitates a ‘biochemical rescue,’ ensuring sufficient enzymatic catalysis to reduce total cyanide levels to safely below the WHO 10 ppm threshold. Units are expressed as *μmol* of HCN released per minute per gram of sample under standardized assay conditions.

## 5. Discussion

### 5.1 Proximate Dynamics: Concentration vs. Fermentative Recovery

The initial processing stage of boiling significantly altered the proximate profile of *Manihot esculenta*. The reduction in moisture from 22.86 g/100g to 20.12 g/100g (Table 1) triggered a concentration effect. This explains the sharp rise in carbohydrate (73.98 g/100g) and energy value (300.68 kcal/mol) observed in the boiled samples. However, fermentation acted as a balancing phase. From 24 to 72 hours, moisture levels showed a slight recovery, which stabilized the caloric density.

Interestingly, protein and lipid contents increased progressively during fermentation, peaking at stage D3. This enhancement is likely due to the secretion of extracellular enzymes by fermenting microorganisms, such as lactic acid bacteria and yeast [8]. These microbes synthesize microbial proteins and lipids, slightly boosting the nutritional density of the tuber. Fiber remained largely stable, suggesting that the structural polysaccharides of the “Farmer’s Pride” variety are resilient to these processing methods [7].

### 5.2 Detoxification Kinetics: The Primacy of Fermentation

Cassava safety is defined by the reduction of cyanogenic glycosides. Fresh tubers contained 98.15 mg/100g of total cyanide, which is dangerously high. While boiling reduced this to 59.02 mg/100g, the levels remained far above the World Health Organization (WHO) safety threshold of 1.0 mg/100g (10 ppm) [6].

The subsequent fermentation stages (D1-D3) were the true drivers of detoxification. By 72 hours (D3), total cyanide plummeted to 0.54 mg/100g (Figure 6), making the tuber internationally safe for consumption. This drastic decline is attributed to the synergistic action of linamarase. Although boiling initially denatures endogenous enzymes, the spike in linamarase activity during fermentation (reaching 8.51 units/g at D3) suggests that microbial enzymes take over the hydrolysis of linamarin and amygdalin [10].

Furthermore, the significant reduction in oxalates (95.2%) and phytates (97.6%) by stage D3 is vital. These anti-nutrients typically bind minerals like calcium, iron and zinc [5]. Their removal suggests that the processed tuber not only becomes safer but also more effective at delivering essential minerals to the consumer.

### 5.3 Vitamin and Antioxidant Resurgence

A critical “nutrient trade-off” was observed during boiling [7]. Vitamin C suffered a 93% loss due to thermal degradation and leaching into the processing water. However, fermentation facilitated a “biochemical rescue.” Vitamin C levels rose from 1.82 mg/100g in the boiled stage to 12.33 mg/100g after 24 hours of fermentation. This increase indicates microbial synthesis of ascorbic acid or the release of bound precursors [2].

The most profound finding for therapeutic research is the transformation of the antioxidant profile. Boiling reduced both TPC and TFC. Conversely, fermentation triggered a massive increase, with TPC reaching 78.32 mg GAE/100g at D3. This nearly doubles the amount found in fresh cassava.

This surge in bioactives is likely due to microbial beta-glucosidases breaking down complex cell wall structures [10]. This process liberates bound phenolics, transforming a simple carbohydrate staple into a bioactive food vehicle. The high antioxidant yield at 72 hours provides a strong biochemical basis for testing these extracts in models of oxidative stress, such as benign prostatic hyperplasia and prostate cancer.

## 6. Conclusion

This study establishes that the safety and therapeutic value of the *Farmer’s Pride (IBA 961632)* cassava variety are strictly dictated by the processing sequence. While a 10-minute boiling period optimizes caloric density through moisture reduction, it fails to mitigate the toxicological risks associated with endogenous cyanogens, leaving levels significantly above the WHO safety threshold.

The most critical finding is the role of fermentation as a “biochemical refinery.” Beyond 48 hours, fermentation serves a dual purpose: it achieves a near-complete detoxification (reducing cyanide to 0.54 mg/100g and facilitates a “biochemical rescue” of the plant’s antioxidant profile. The significant surge in TPC and TFC at the 72-hour (D3) stage suggests that microbial activity liberates bound bioactives from the plant cell wall. Consequently, the sequential application of boiling followed by extended fermentation transforms this staple tuber from a simple energy source into a safe, bioactive food vehicle with high potential for therapeutic intervention in oxidative stress-mediated pathologies.

## 6. Recommendations

Based on the evidence, the following multi-level recommendations are proposed:

- **For Food Safety and Household Processing:** Public health campaigns should discourage the consumption of boiled-only cassava. A standardized 72-hour fermentation protocol must be adopted to ensure cyanide levels drop below the 10 ppm safety limit. The use of uncovered vessels is mandatory to facilitate the volatilization of hydrogen cyanide (HCN) gas.
- **For Agricultural and Public Health Policy:** The *Farmer’s Pride (IBA 961632)* variety should be integrated into biofortification and food security programs in Nigeria. Its robust metabolic response to fermentation makes it a superior candidate for producing nutrient-dense, safe food products for low-income populations.
- **For Clinical and Translational Research:** 1. **Analytical Profiling:** Future studies should utilize High-Performance Liquid Chromatography (HPLC) or LC-MS/MS to characterize the specific phenolic profile (e.g., gallic acid, ferulic acid, or catechin) released during the D3 stage. 2. **Pharmacological Evaluation:** Given the observed peak in antioxidant capacity, the D3 extract should be prioritized for *in vivo* pharmacological screening. Specifically, its efficacy in modulating androgen-induced oxidative stress and pro-inflammatory cytokines should be evaluated in models of Benign Prostatic Hyperplasia (BPH) and prostate cancer

## Acknowledgments

The authors wish to thank the **National Root Crops Research Institute (NRCRI), Umudike, Abia State**, for the botanical validation and variety coding of the *Farmer’s Pride* (IBA 961632) cassava variety. We also acknowledge the laboratory staff of the **Department of Biochemistry, University of Uyo**, for providing access to the spectrophotometric facilities used for the assays except the enzyme activity assays.

## Ethics Statement

The study was conducted in accordance with local and institutional guidelines for laboratory research. Ethical approval was obtained from the Faculty of Basic Medical Sciences Research Ethics Committee (FBMSREC), University of Uyo, Nigeria (Approval No: **UU FBMSREC 2025 020**). No human or animal subjects were used in this biochemical analysis; however, all plant materials were handled according to institutional botanical safety standards.

## Data Accessibility

The datasets supporting the conclusions of this article, including raw proximate analysis values, anti-nutrient concentrations, and spectrophotometric linamarase activity data, are available on Figshare at https://figshare.com/s/ac2942617ec2e4fc1e0c?file=62090488.

## Authors’ Contributions

In accordance with the CRediT (Contributor Roles Taxonomy) framework, the contributions are as follows:

- **G.E. Bassey:** Conceptualization, Methodology, Investigation, Data Curation, Formal Analysis, Writing –Original Draft, and Project Administration, Resources, Visualization
- **E.O. Jimmy:** Supervision, Conceptualization, Resources, Writing –Review & Editing.
- **H.T. Olatunbosun:** Co-supervision, Validation, Data Curation, Writing –Review & Editing.

All authors have given final approval for the version to be published and agree to be accountable for all aspects of the work.

